# Gestures Of Hunger: Uncovering Intentional Gestural Communication In Free-Ranging Hanuman Langurs

**DOI:** 10.1101/2023.09.07.552163

**Authors:** Dishari Dasgupta, Arnab Banerjee, Akash Dutta, Shohini Mitra, Debolina Banerjee, Rikita Karar, Srijita Karmakar, Aparajita Bhattacharya, Swastika Ghosh, Pritha Bhattacharjee, Manabi Paul

## Abstract

Contrary to previous beliefs, intentional gestural communication (IGC) is not exclusive to the hominoid lineage but is also present in other non-human primates. Here, we report the presence of IGC among free-ranging Hanuman langur troop in Dakshineswar, West Bengal, India. These langurs exhibit a food-requesting behaviour wherein they use several gestures to communicate with the humans nearby. Moreover, they can also assess the recipient’s mental state and persistently check if the signal (food request) has been received, waiting until they receive the desired food item. We have identified eight begging gestures used by langurs of all ages, except infants. The most common gesture is by holding cloth (BGc), but provocation-initiated begging (BGpi) and begging by embracing legs (BGe) efficiently direct these events to its success. The frequency of successful begging events is higher in the evening due to increased human interactions. Our findings suggest that ontogenetic ritualization might be at play here among these troop members as this gestural communication has been learned through imitation and reinforced by the reward of receiving food. Moreover, these successful begging events serve as an effective foraging strategy for urban-adapted langurs, allowing them to acquire high-calorie processed food items within a human-modified urban ecosystem.

## INTRODUCTION

The evolution of language has been a subject of interest for researchers since the eighteenth century. Language is often considered one of the defining features of human evolution (Corballis,2017). Besides language, human children habitually use gestures as a primary mode of communication and eventually acquire the ability to speak (Meguerditchian et.al.,2011). However, research on non-human primates (NHPs) has revealed that they also possess a range of communication abilities, including intentional gestural communication (IGC). Initially, it was suggested that language evolved from gestures, known as the ‘gesture-first hypothesis’ (Hewes,1973). In support of this theory, researchers taught American sign language to a captive female chimpanzee, who successfully started communicating after 22 months, by using a combination of various gestures (Gardner & Gardner,1969). Studies conducted on captive Gorillas also confirm that they can exhibit IGC wherein they take into account the attentional state of the receiver, a hallmark of ‘true communication’ (Pika et al.,2003).

Intentional communication refers to communication wherein the sender and receiver take into account each other’s mental states and behaves accordingly. It can be divided into zero-order and first-order intentionality. Zero-order intentionality occurs when the sender does not consider the recipient’s thoughts and does not realize how their actions affect the recipient’s mental state. Whereas, in first-order intentionality, the mental state of the recipient is taken into account by the sender and the sender also recognizes the effect of its’ behaviour on the mental state of the recipient (Dennett 1983; Call and Tomasello 2007).

Studies conducted on several ape species like Bonobos, Chimpanzees, Gorilla, Siamangs and Orangutans have confirmed the presence of IGC in NHPs. (Chimpanzee - Goodall,1968; Tomasello et al., 1994; Roberts et al., 2012; Gorilla - Pika et al., 2003; Genty et al. 2009; Bonobos - Genty et al., 2015; Graham et al., 2017; Orangutans - Liebal et al., 2006; Siamangs – Liebal et al.,2004). Additionally, studies on Collared mangabey, Tufted capuchins, Common squirrel monkey, Wild spider monkey, Olive baboon, Tonkean macaque and Rhesus macaque, have also demonstrated IGC between monkeys and humans (Schell,2022; Red-capped mangabays - Aychet et al. 2020; Tufted capuchin monkeys - Hattori et al.2010;; Squirrel monkeys - Anderson et al. 2010; Mangabeys - Maille et al. 2012; Olive Baboons - Molesti et al. 2020; rhesus macaque - Deshpande et.al.,2018; Canteloup et al. 2015; Bonnet macaque - Gupta and Sinha,2019; Wild spider monkey - Larenas et.al.,2023). All these studies indicated that IGC is not unique to humans, but is also present in NHPs. Hobaiter and Byrne (2011) found that chimpanzees can use different gestures, such as arm raises, head nods, and hand claps, to establish meaningful communication, such as greeting, threat, and invitation. Additionally, Gupta and Sinha (2019) observed IGC consisting of visual and tactile gestures among the wild bonnet macaques of Bandipur National Park, India. All these studies suggest that NHPs possess the ability to use various gestures to intentionally communicate with their conspecifics. Furthermore, Cartmill and Byrne (2007) observed captive orangutans to use a wide range of intentional as well as goal-directed gestures for communicating with their caregivers, which indicates the presence of a high level of cognitive complexity in their communication. Pika and her colleagues (2003) found ‘ontogenetic ritualization’ as the underlying mechanism for learning such gestural communication in NHPs. The ‘ontogenetic ritualization hypothesis’ (ORH) and the ‘phylogenetic ritualization hypothesis’ (PRH) are two competing theories that explain how non-human primates learn and use gestural communication, thereby continuing debate and controversy. While the former emphasizes the role of social learning and experience in the development of gestural communication, the latter focuses on the innate biological predisposition for gestural communication (Call & Tomasello, 2007; Hobaiter & Byrne, 2011).

Apart from acknowledging the attentional states of the recipient, another hallmark of human language is the ability to use different gestures for the same goal and/or use the same gesture for different goals, known as means-end dissociation (Bruner, 1981; Call and Tomasello, 2007; Pollick and de Waal, 2007). While IGC with goal persistence and means-end dissociation has been reported in some NHPs, most of them were conducted in artificially constrained environments, leaving a lacuna of comprehensive studies on the catarrhine lineage (Schell et al., 2022; Molesti et al.,2020). To address this gap in our understanding of language evolution in primates, we conducted a study on a free-ranging Hanuman langur troop residing in a temple area of West Bengal, India.

Hanuman langurs (*Semnopithecus entellus*) are folivorous colobines, which usually locomote as unimale-multifemale troops and all-male bands (Konecna et al.,2008). Even though there seems to be confusion regarding Hanuman langurs being arboreal or terrestrial, our study troop is mostly terrestrial as they spend considerable amount of their time interacting with humans (Dutta et.al., 2023; Dasgupta et.al., 2021). To the best of our knowledge, this study has given the first systematic evidence of the presence of intentional gestural communication (IGC) with goal persistence and means-end dissociation in the highly provisioned free-ranging langur troops residing in Dakshineswar, West Bengal, India. Besides identifying the probable factors contributing to successful communication using gestures, our results have also revealed how these langurs used several different gestures to direct devotees to provide food to them and persisted until they received the preferred food item. Additionally, we have also examined whether such gestures are species typical or did they learn by imitating their fellow troop members. Altogether, this study suggests that pre-existing communication abilities in other primates, including the catarrhine lineage, may have laid the foundation for the evolution of human language (Schell et al., 2022).

## METHODS

### Study area and study troop

We conducted our field study between January, 2019 and April, 2021 on free-ranging Hanuman langur (*Semnopithecus entellus*) troops in various parts of West Bengal, India. Three distinct troops were identified for long-term observations based on the degree of anthropomorphic interactions received by them – Dakshineswar (22.6573° N, 88.3624° E), Nangi (22.4973° N, 88.2214° E), and Sarenga, Nalpur (22.5307° N, 88.1840° E). Observers recorded the location details of each langur troop along with respective troop size and behaviours (both intra- and inter-specific interactions). *Ad libitum* data were used to classify langur’s life stages into four distinct categories based on their physical and behavioural characteristics; 1. Infant-dark fur colour and fully dependent on adults for their movement and feeding; 2. Juvenile-light fur colour similar to adults but smaller in size and partially dependent on adults for their movement and feeding; 3. Subadult-fully independent of adults but yet to attain sexual maturity, body size is typically in between that of juveniles and adults; and 4. Adult-fully grown, independent individual who is sexually mature. Besides, an ethogram was generated wherein two to three letter codes were assigned to distinct recorded behaviours followed by their detailed description. With maximum anthropomorphic interaction being recorded in Dakshineswar area (Dasgupta et al., manuscript in preparation), ‘begging’ was only observed among the troop residing in Dakshineswar in contrast to Nangi and Nalpur, where we did not observe this behaviour.. Hence, we chose the langur troop residing in the Dakshineswar temple area as the focal group for recording the begging behaviour in detail.

### Data collection

The langur troop (DG) residing in Dakshineswar temple area was observed in three different sessions of the day; morning (M, 9:00 AM – 12:00 PM), afternoon (N, 12:00 PM-15:00 PM) and evening (E, 15:00 PM – 18:00 PM). We noted langur behaviours of the focal group, DG, by using a combination of *instantaneous scans* and *all occurrence sessions* (AOS) (Altmann, 1974). Furthermore, all instances of begging behaviours were recorded along with the details of initiator, recipient, types of gestures being used by the langurs, whether the act is successful or unsuccessful, location and time. Simultaneously, all these behaviours were also recorded in SONY HDR-CX405 digital video camera. For each session, human footfall and number of langurs present in the area were also recorded.

Here, begging is noted as a gestural communication displayed by the langur who requests humans for food offerings. This event starts when a free-ranging Hanuman langur spots a human being close to vendors selling food items that are consumed by langurs. Langurs display their interest in the food available on the vendor’s cart by extending their hand forward, followed by several postures by using their forelimbs and hindlimbs, to request the human being (buyer) for food provisioning. For a given begging instance, the full duration from asking for food to final consumption by the langur was recorded by the observer and was considered a ‘successful’ begging event. If the langurs refused to accept the food item given by the human, food was considered to be rejected. Moreover, when a human refused to comply, begging event was considered to be ‘unsuccessful’ and the langur’s reaction (if any) was noted down. Here, we categorized the begging behaviours into seven distinct categories based on the posture used by the langur– i) BGp – Bipedal begging, ii) BGq – Quadrupedal begging, iii) BGe – Begging by embracing legs, iv) BGc – Begging by holding cloth, v)BGh – Begging by holding hands vi) BGa – Begging with aggression, vii) PB-Passive begging and viii) BGpi - Provocation initiated begging (Figure 1).

**Figure 1:**
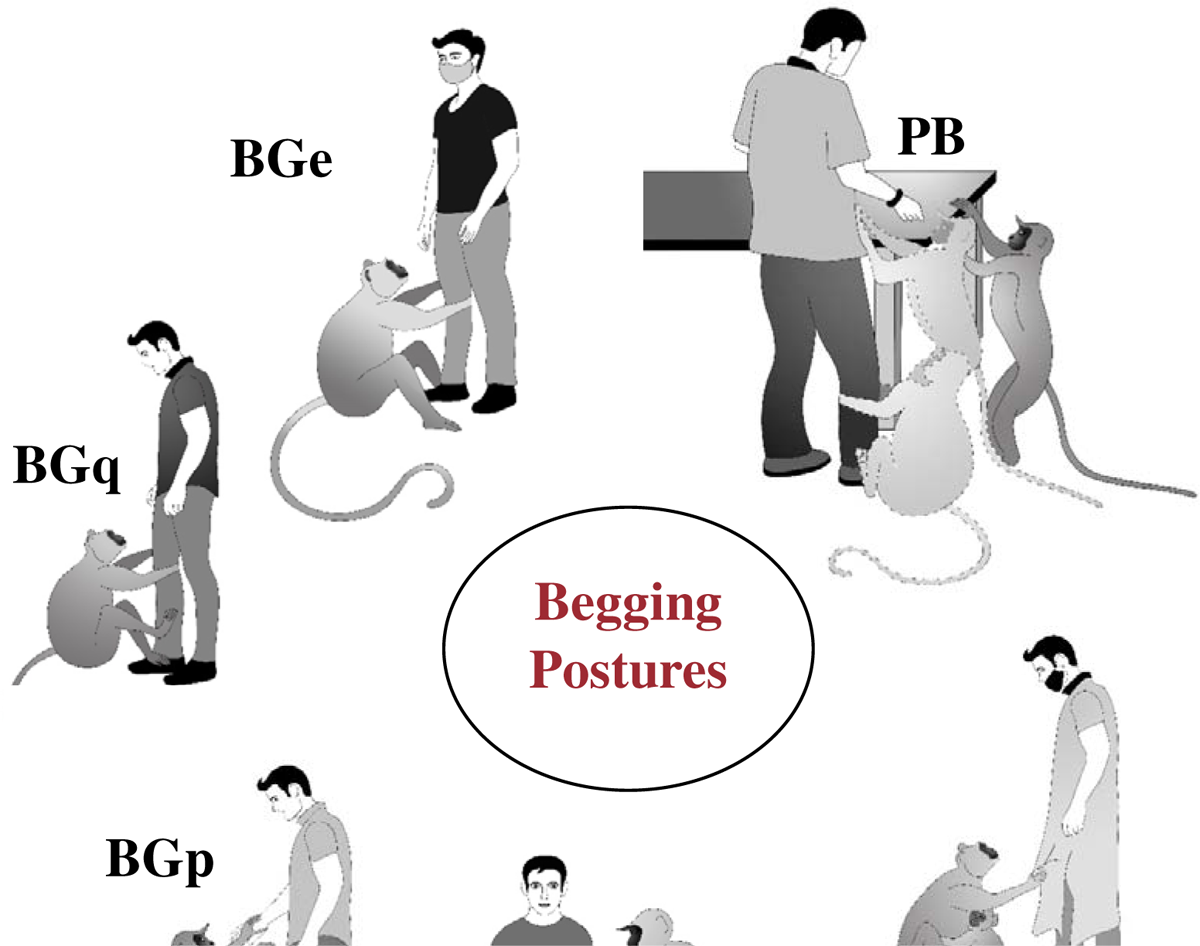
Digital art of begging postures used by the free-ranging langurs in Dakshineswar.

### Data analysis

To determine the factors affecting both successful and unsuccessful begging events, correlation analysis was performed with respect to different begging modes, age and sex of langur, and begging sequence. Here begging sequence has been denoted as ‘offer’ sequence by the signal recipient human being (who receive begging behaviours from langurs) where *first food offering event* is Offer 1, second food offering event is Offer 2 and so on. The observational events were analysed using both categorical and numerical predictive modelling approaches, following Principal Component Analysis (identifying most important factors) and Collinearity tests (eliminate multicollinearity among factors). Our data set was tested for collinearity using VIF or variance inflation factor extraction (Farrar and Glauber, 1967; Thompson et al., 2017) following the algorithm as presented below:

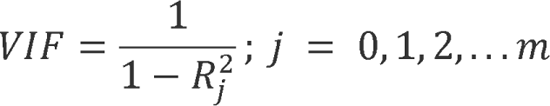

where, *R^2^_j_* represents the unadjusted coefficient of determination for regressing the ***i***th independent variable on the remaining ones. As a rule of thumb, VIF > 5 represents multicollinearity i.e., high correlation among themselves. But for the present case, since some of the variables with the above criteria were instinctively deemed significant, they were not omitted from the final analyses (BGp, BGq, BGe, Bgc, Adult Female, and Offer 1).

### Categorical prediction using Random Forest Models

Random Forest (RF) is a predictive tool first proposed by (Ho, 1995) and later popularized by (Breiman, 2001). These are ensemble classifiers operating by constructing multiple decision trees during model training. In this study, our dataset was categorized into four classes (on the basis of success rate of begging, i.e., number of events yielding successful outcome): low (1-10), moderate (11-20), high (21-30), and extremely high (31 and above). RF models were used to understand and predict these categorical groupings based on our explanatory variables like different begging modes, age and sex of langur, begging sequence, and human-langur footfall.

### Begging prediction using Artificial Neural Networks

While RF models are useful in categorical prediction, Artificial Neural Networks (ANN) are effective tool for analysing the complex non-linear relationships in apparently non-related data. The architecture of an ANN represents the biological neurones of an organism. They can read, interpret, understand inputs, and provide feedbacks based on that. The ANN architecture consists of input layer (with predictors), hidden layer (response and calculation layer) and output layer (predicted outcome) (Gallant, 1993; Ham and Kostanic, 2000; Smith, 1993).

In the present scenario, there are 52 nodes in the hidden layer with 17 input variables and one output. A total of 9225 models were evaluated using only a single hidden layer. The response variable (output) is the successful event count (Succ). The models were then evaluated for prediction and subsequently root mean square error (RMSE) has also been calculated for model validation based on the predicted values (Figure 2).

**Figure 2:**
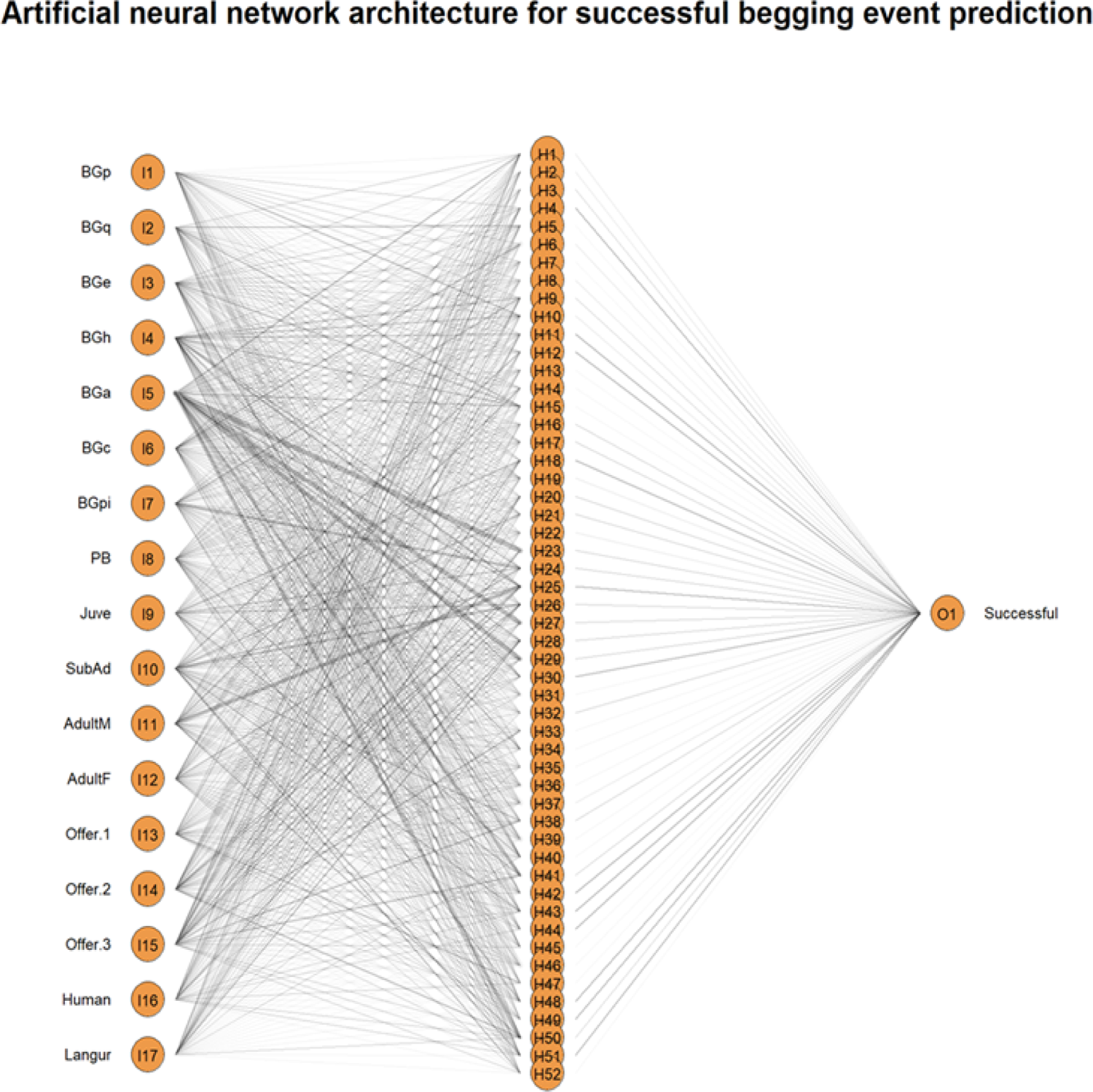
Architecture of Artificial Neural Network. There are 52 nodes in the hidden layer with 17 input variables and 1 output. Total number of models evaluated = 9225 for single hidden layer, and 3.9852^{4} for 2 hidden layers, with success counts (Succ) as output.

### Ethical note

No langurs were harmed during this work. All work reported here was purely observation-based and did not involve direct handling of langurs in any manner, therefore, was in accordance with approved guidelines of animal rights regulations of the Government of India. The research reported in this paper was sanctioned by DST-INSPIRE, Government of India (approval number: DST/INSPIRE/04/2018/001287, dated 24th July 2018), and was also notified to the Principal Chief Conservator of Forests (PCCF), West Bengal, India (Memo No. 869/WL/4R-43/2021, Date: 07.04.2021).

## RESULTS

### Successful Begging Event Data

The mean and median of successful events are 11.13 and 7.00 respectively. However, the highest value (65) suggests that there are outliers in the data-set (Figure 3). Outliers have been discarded and data has been standardized for further predictive modelling.

**Figure 3:**
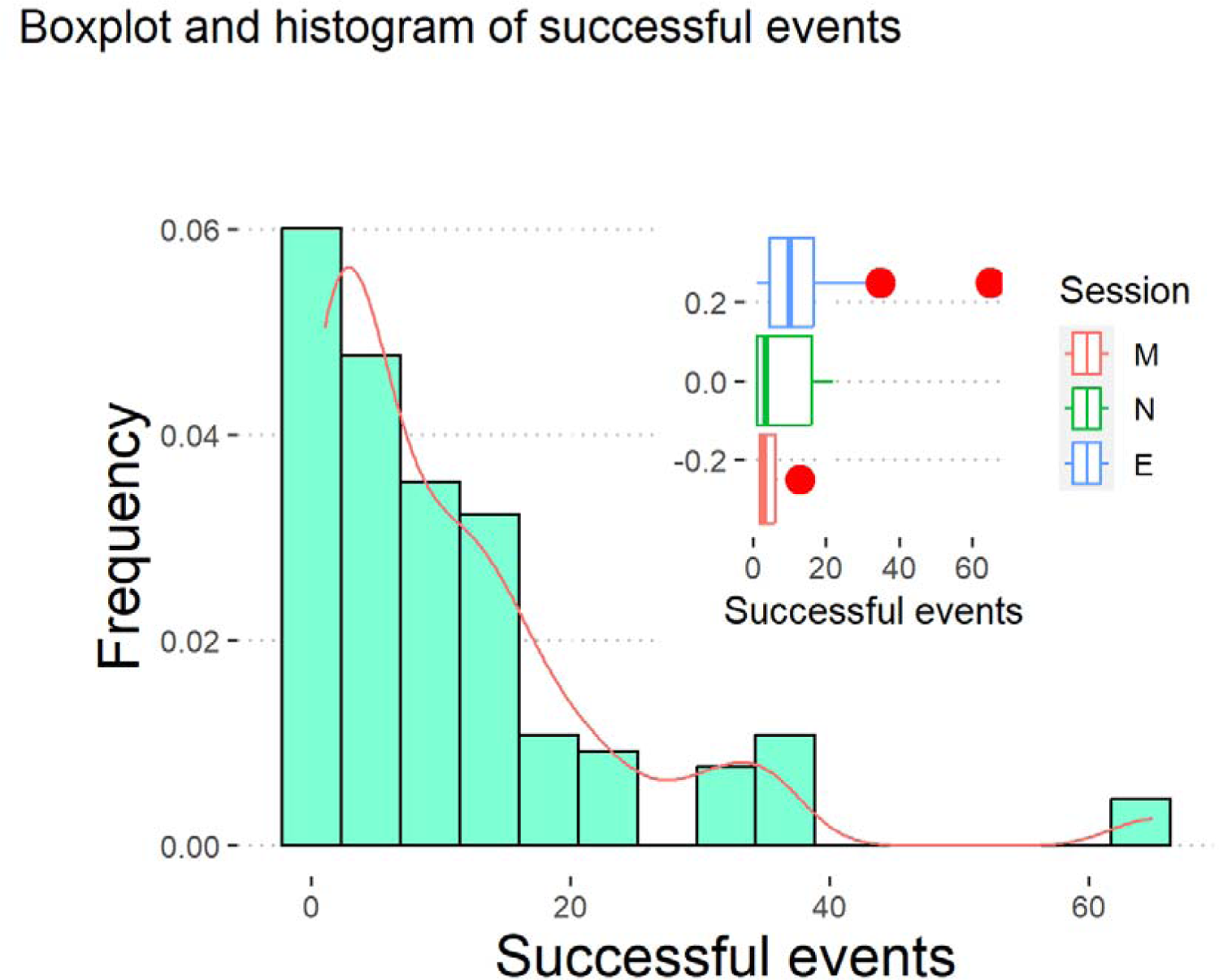
Box plot and histogram. The mean and median of successful events are 11.13 and 7.00 respectively. However, the highest value (65) suggests that there are outliers in the data-set.

### Factors affecting successful begging

Successful begging events and unsuccessful events had varied levels of correlation with the modes of begging. The most efficient methods of begging are BGpi (*r* = 0.801) followed closely by BGe (*r* = 0.772), BGc (*r* = 0.729) and PB (*r*=0.691). Whereas BGc (*r* = 0.777) leads to most unsuccessful begging, followed by BGq (*r* = 0.688) (Correlation: Figure 4a). While all the adults have most success in begging (*r* = 0.965), adult females have high correlation with both successful (*r* =0.963) and unsuccessful begging event (*r* = 0.758) (Figure 4b). For offer sequence, highest correlation with successful begging event was observed for the initial begging sequence (*r* = 0.988). Interestingly, correlation between successful begging event and begging sequence decreased as the begging sequence proceeded further to sequence two (*r* = 0.325) and three (*r* = 0.061) (Figure 4c).

**Figure 4:**
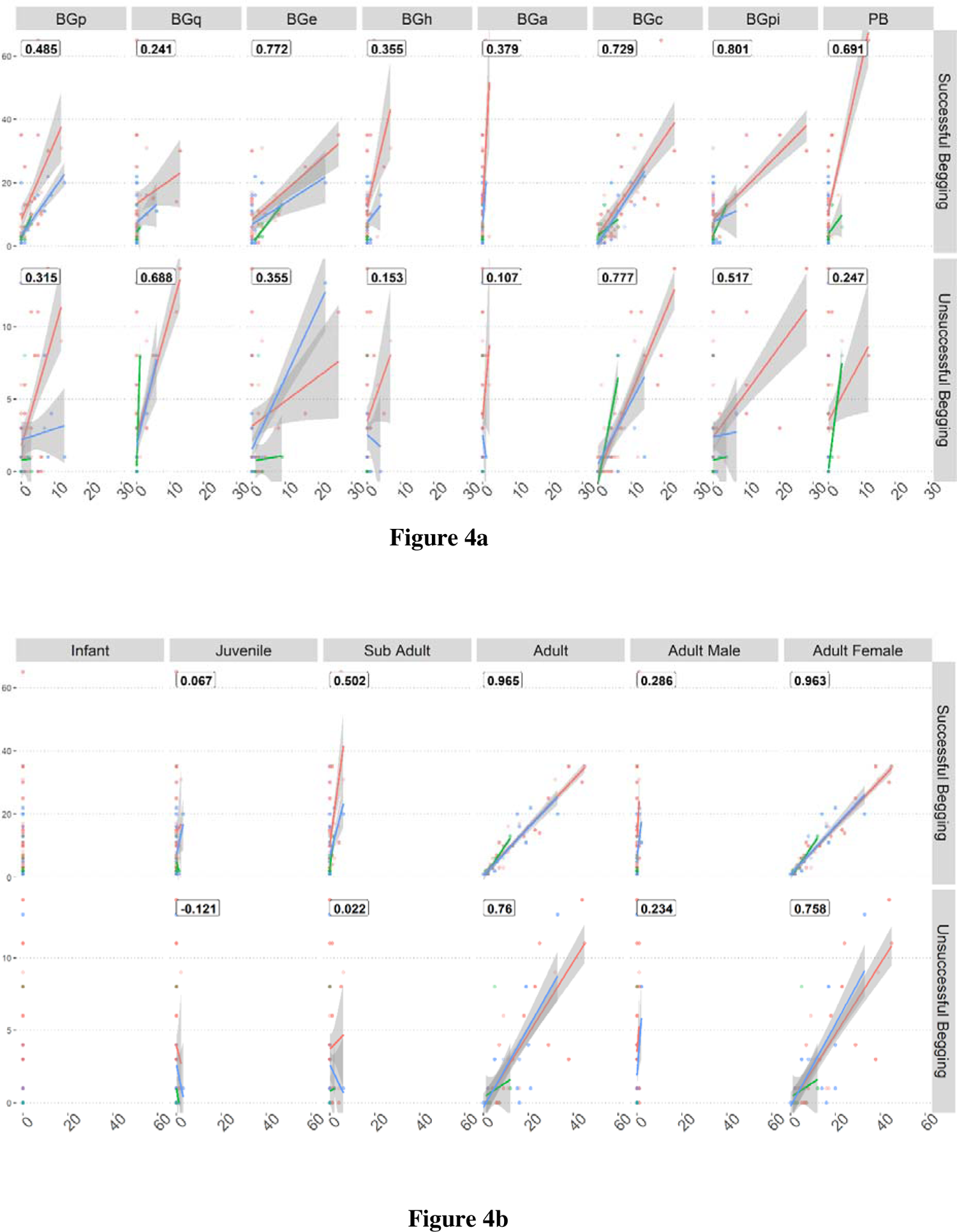
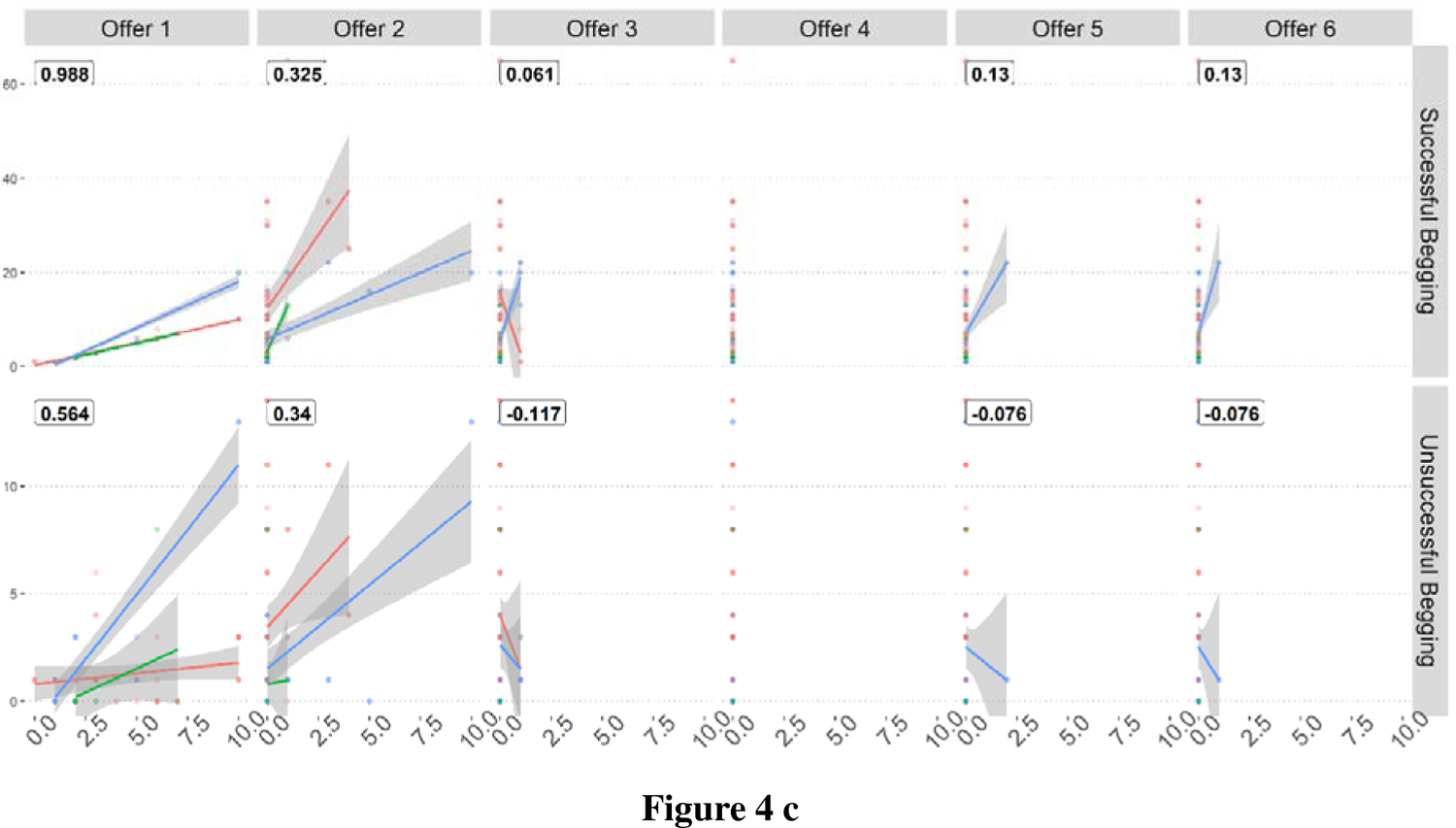
Correlation plots. **4a)** Correlation among begging success and modes of begging. The most efficient methods of begging are BGpi (*r* =0.801) followed closely by BGe (*r* =0.772), BGc (*r* =0.729) and PB (*r* = 0.691). Whereas BGc (*r* =0.777) leads to most unsuccessful begging, followed by BGq (*r* =0.688). **4b)** Correlation among begging success and age / sex of langur. All the adults have most success in begging (*r* = 0.965), adult females have high correlation with both successful (*r* = 0.963) and unsuccessful begging event (*r* = 0.758). **4c)** Correlation among begging success and offerings. highest correlation with successful begging event was observed for the initial begging sequence (*r* = 0.988). But, correlation between successful begging event and begging sequence decreased as the begging sequence proceeded further to sequence two(*r* = 0.325) and three(*r* = 0.061).

Results of PCA revealed that most of the variability in the experimental observations can be explained through PC1(69.61%), PC2 (13.33%) and subsequently another 6.35% by PC3 making a cumulative total of 89.29% explained variance over the first three components (Table 1). Along PC1, adult female (0.659) and BGc (0.203) has the highest and lowest value respectively, suggesting that these two variables have higher impact in determining the success rate of a begging event. Along PC2, human (0.903) and BGe (−0.246) has the highest and lowest value respectively suggesting that these two variables have higher impact in bringing about a successful begging event (Table 1). From the PCA biplot it can be observed that even though begging occurs across all three sessions – morning, noon and evening, most of it is observed in the evening (Figure 5).

**Table 1:**
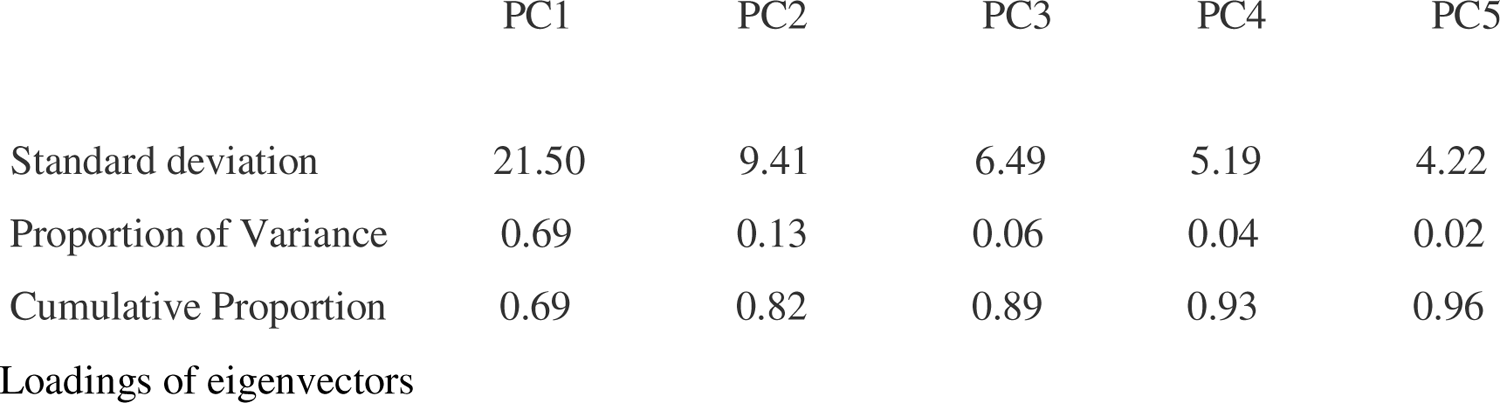
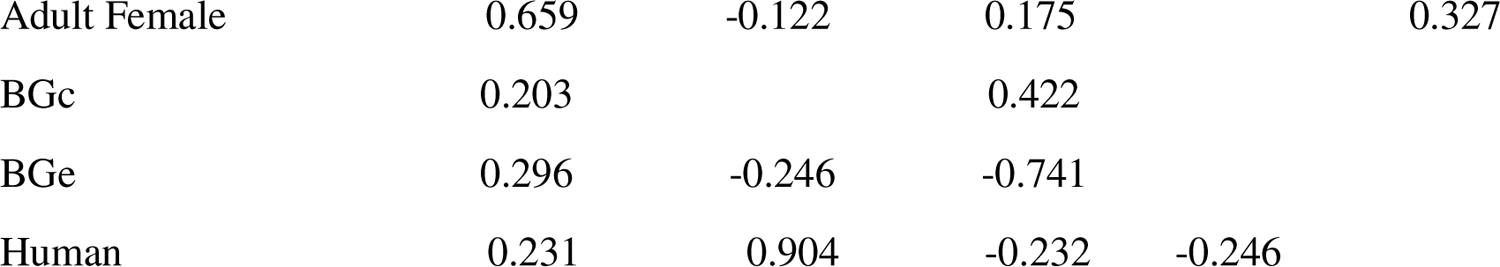
Principal Component Analysis. Results of PCA show that most of the variability in the experimental observations can be explained through PC1 (69.61%), PC2 (13.33%) and subsequently by PC3 (6.35%) making a cumulative total of 89.29% explained variance over the first three components. Along PC1, adult female (0.659) and BGc (0.203) has the highest and lowest value respectively. Along PC2, human (0.903) and BGe (−0.246) has the highest and lowest value respectively.

**Figure 5:**
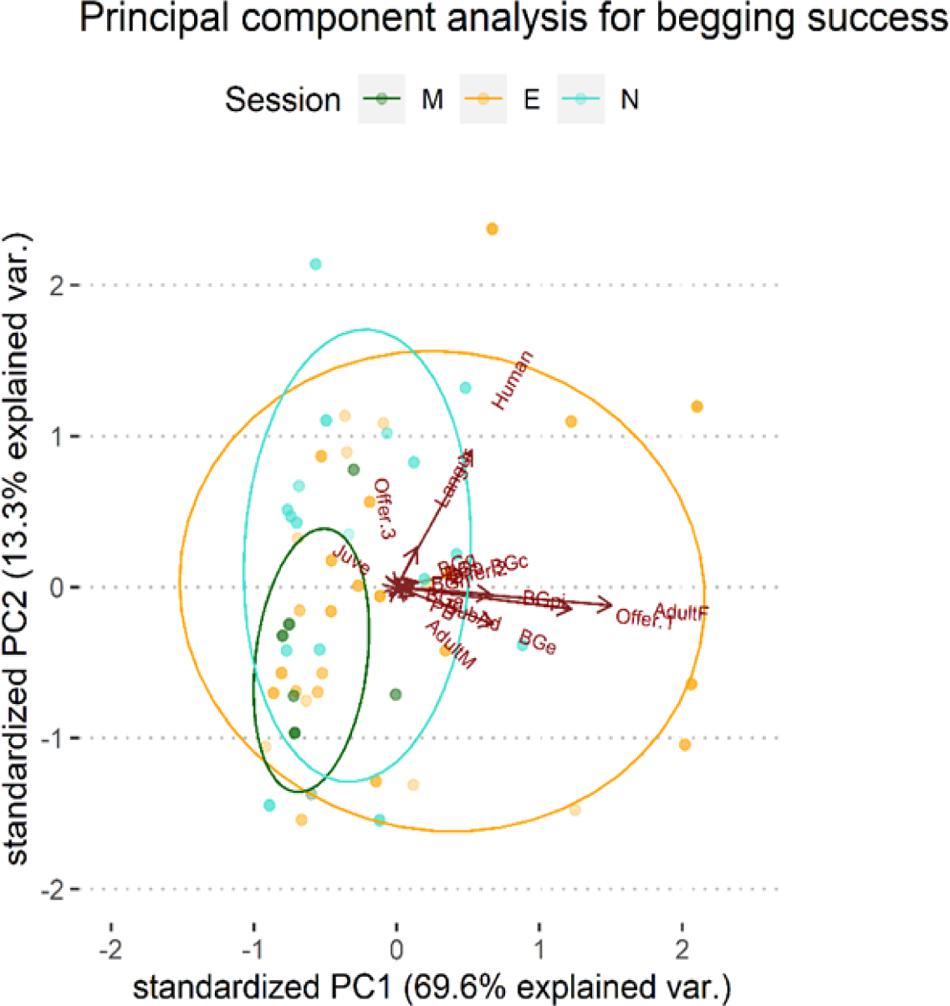
PCA Biplot. PCA biplot shows that even though begging occurs across all three sessions – morning, noon and evening, most of it is observed in the evening.

### Collinearity

Often the independent predictor variables have high correlation among themselves which gives rise to a situation that generate false predictive accuracies. In the present case, after VIF extraction, we omitted the following predictors: Infant, Adult,Offers 4∼6, Inactive, Solitary, Pair, Group, Forage, Human offer, Eating.

### Random Forest Model

The error rates are stabilized after around 200 trees (approx visualization). The OOB or *out-of-bag* error line shows a decrease till about 270 trees beyond which it plateaus without any further reduction. The value of OOB error = 0.88% denotes that there is a chance of only 4.4 trees out of 500 trees trained, giving an erroneous result from the predicted model. We observed that the *first food offering event* and the involvement of *adult female* langurs are key to better begging success (Figure 6). From the results of the confusion matrix analysis, we can see that our RF model is highly efficient in predicting success rates, with about 96.43% accuracy (Table 2). There is only one misclassification in the predicted data (moderate classified as extremely high - hence sensitivity of class prediction of moderate is 90%). Our predictive model has accuracies between 81.65% and 99.91% at 95% level of confidence. Thus, we can now predict the different classes of begging success rates. For a more accurate (and numerical) prediction, we have used artificial neural networks.

**Figure 6:**
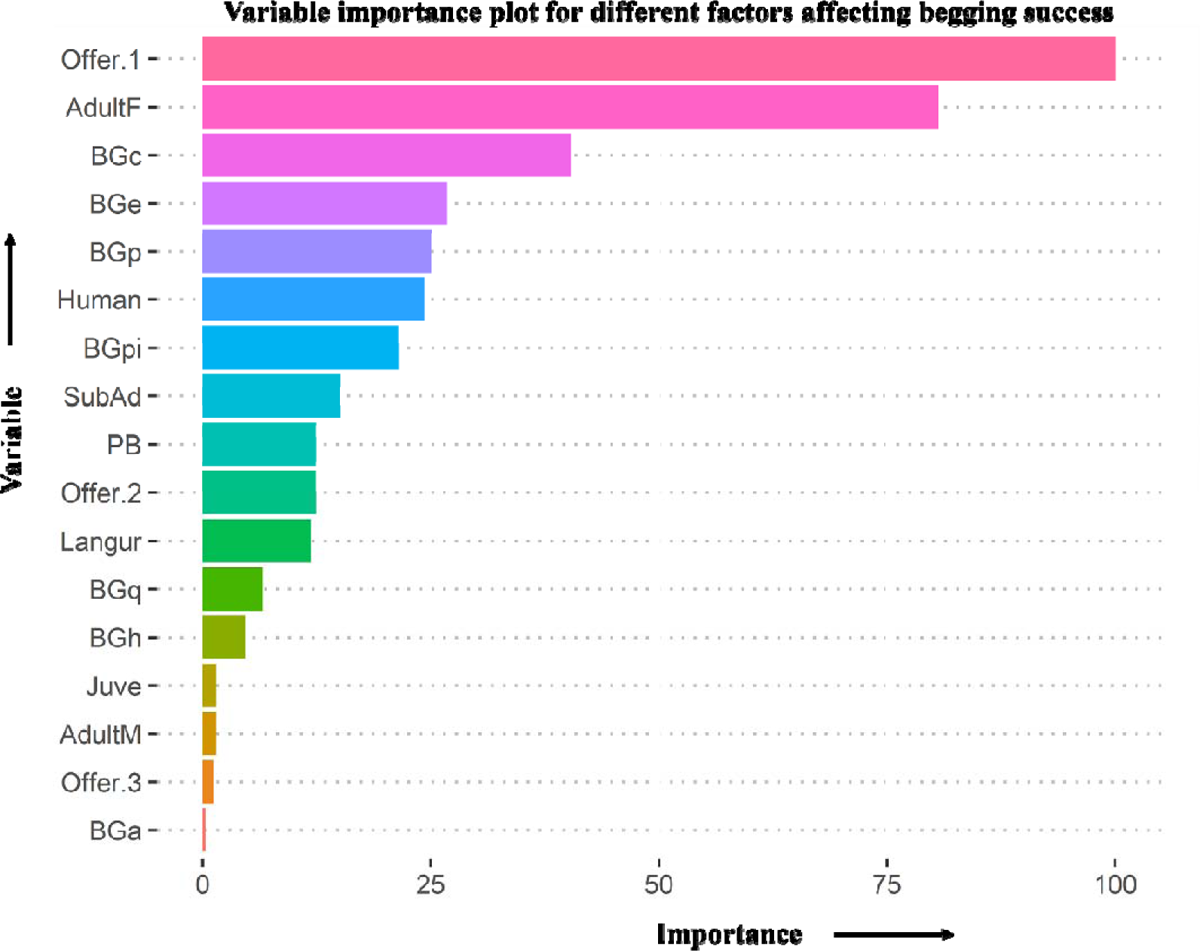
Variable importance plot for different factors affecting begging success. The first food offering event and the involvement of adult female langurs have highest importance in begging success.

**Table 2:**
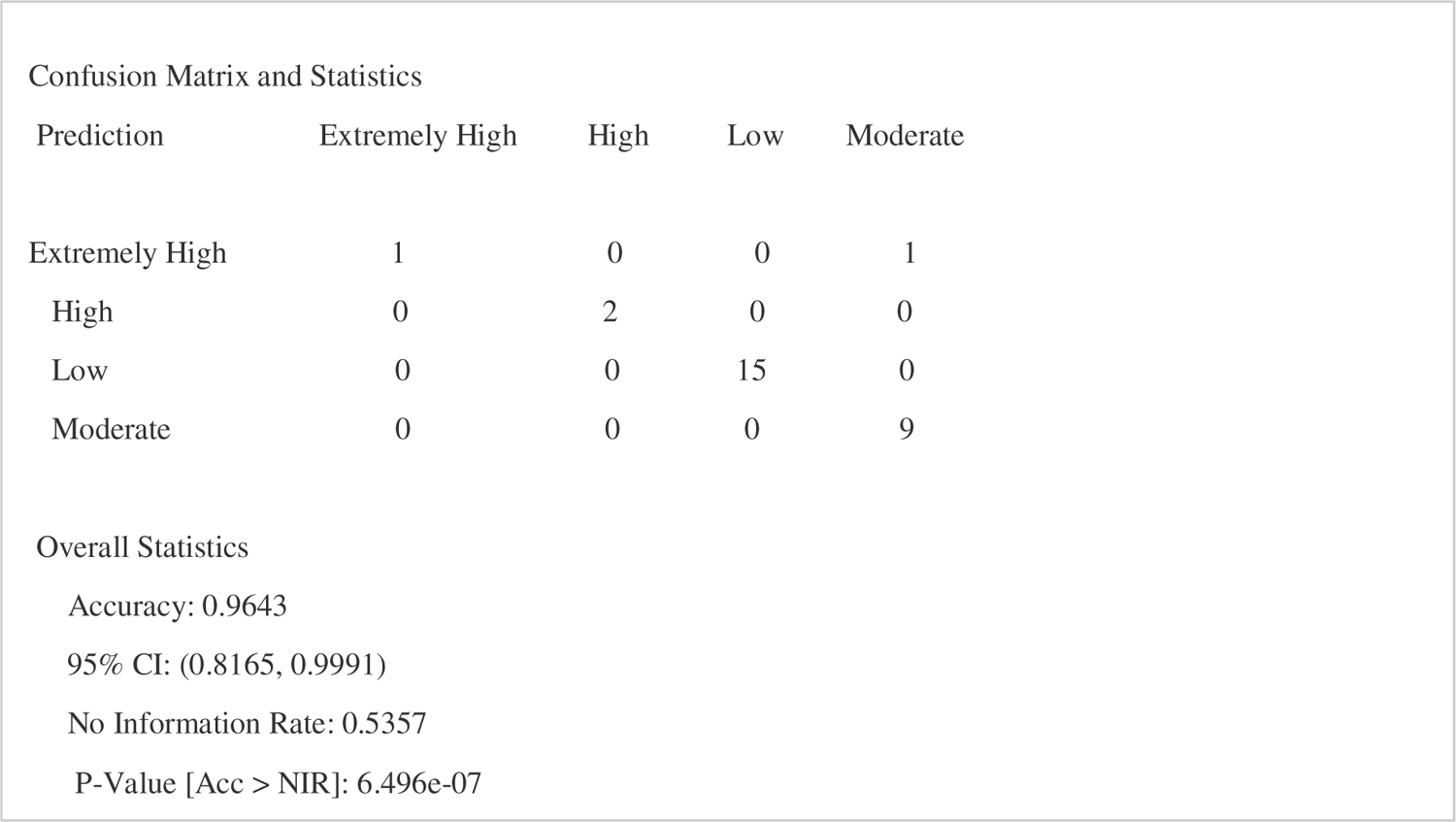
Confusion matrix. Confusion matrix analysis shows that, Random Forest model is highly efficient in predicting success rates, with about 96.43% accuracy. There is only 1 mis-classifications in the predicted data (moderate classified as extremely high - hence sensitivity of class prediction of moderate is 90%). Our predictive model has accuracies between 81.65% and 99.91% at 95% level of confidence.

### Artificial Neural Network

The RMSE for the model is calculated to be 0.259 and from the plot we can see that our observed and predictive value is very close (Figure 7). The sensitivity plot (Figure 8 a.b.c) shows that adult female, begging sequence 1 (Offer 1) and provocation-initiated begging are the most efficient descriptor for successful begging events. These results are in agreement with the previous observations of the RF model and correlation analysis.

**Figure 7:**
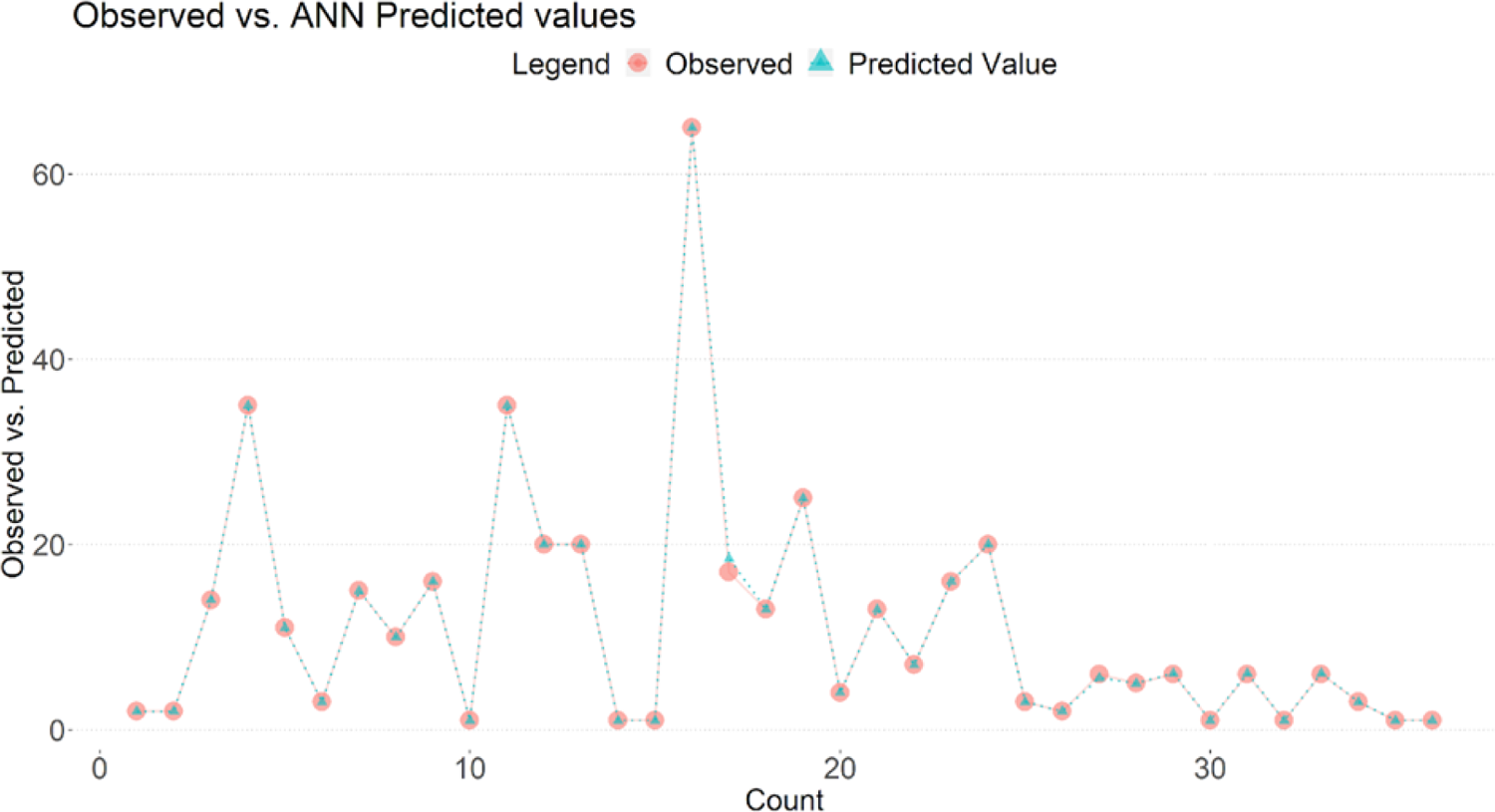
Graphical representation of Observed vs ANN Predicted values. From the plot we can see that our observed and predictive value is very close.

**Figure 8:**
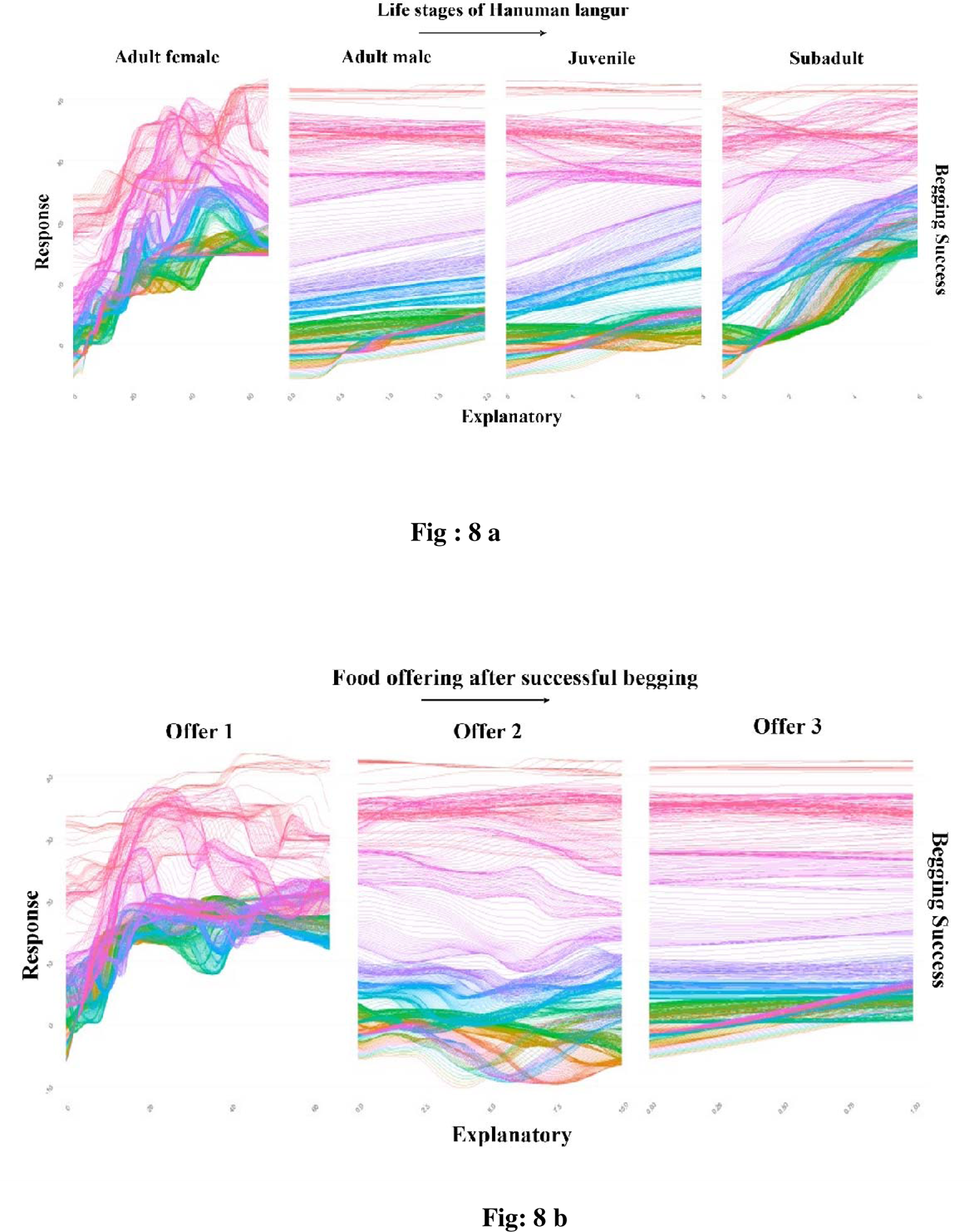
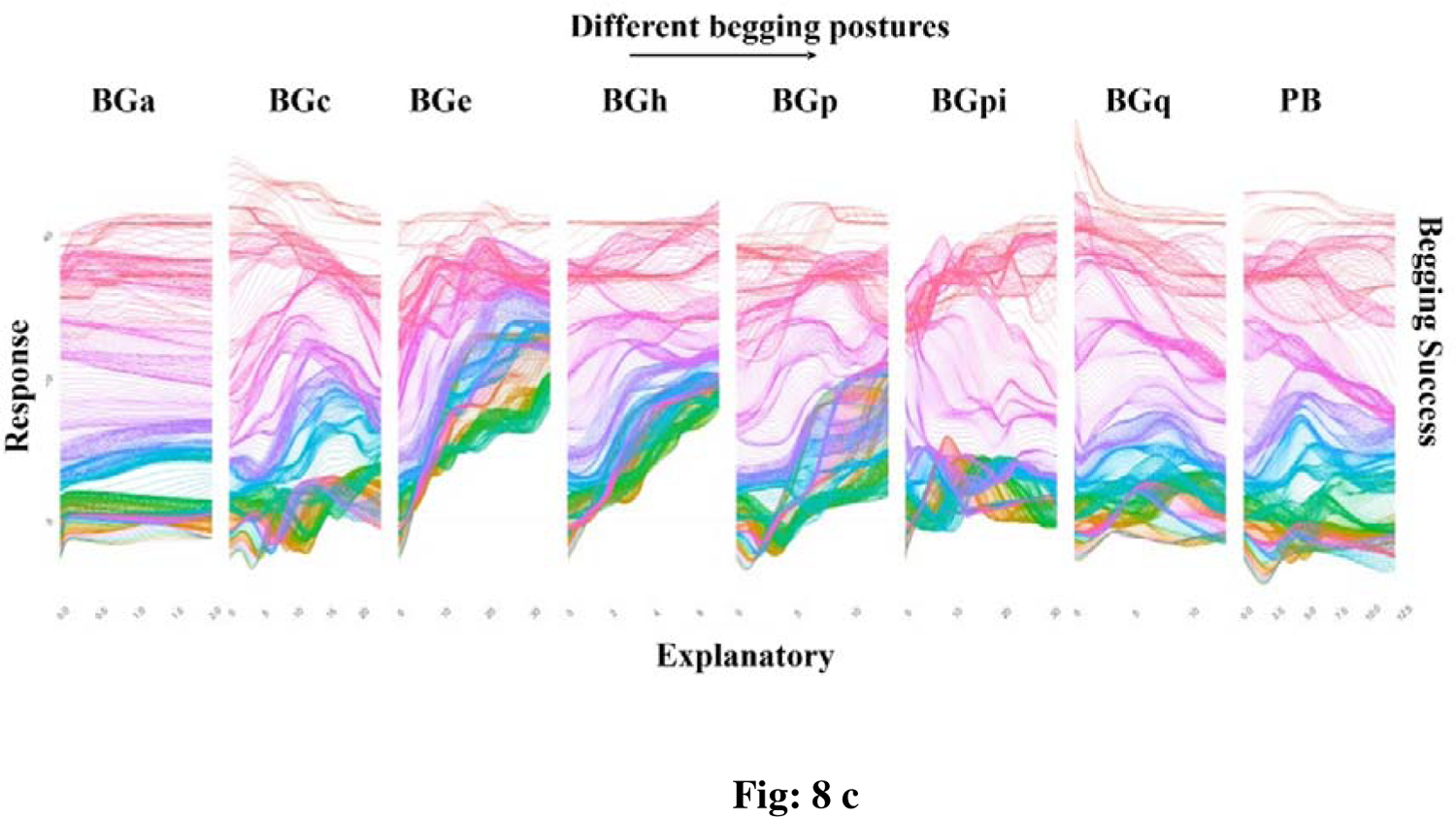
The following plots show the various sensitivity of the explanatory variables towards the final dependent evaluation. **8a)** Sensitivity towards the age and sex of Langur. This plot shows that adult female is the most efficient descriptor for successful events over others. **8b)** Sensitivity towards offer sequence, this plot reveals that offer sequence 1 is the most effective and sensitive parameter. **8c)** Sensitivity towards begging modes wherein we can confirm that provocation-initiated begging is the most efficient contributor towards successful events.

## Discussion

This study delineates the presence of first-order intentional gestural communication in free-ranging Hanuman langurs residing in Dakshineswar, West Bengal, India. Here, these langurs not only assess the mental state of the receiver (human in this case), but also keeps checking on the human to see whether the intended recipient has in fact received the given signal (of food request by langur), and waited until the human has given the food item of its choice (sign of goal persistence). Besides, these langurs used different types of gestures (various begging postures) to convey one single message – “give me food” (means-end dissociation), which together with ‘goal-persistence’, fulfill all the three criteria of a true intentional gestural communication (IGC). Earlier studies reported that the point of origin with regard to human language was present only in hominoid lineage in the form of intentional and flexible manual gestures (Call and Tomasello, 2007). But our results contradict such claim. Rather, it is in line with recent findings which reports the presence of IGC in the platyrrhine and catarrhine lineage as well consisting of bonnet macaques, olive baboons, tonkean macaques, spider monkeys and red-capped mangabeys (Cateloup et.al.,2015; Gupta and Sinha,2019; Molesti et.al.,2020; Schel et.al.,2022; Larenas et.al.,2023). Our study troop uses a total of eight different gestures for begging, all of which are shown by both the sexes and across all life stages, barring infant. These gestures are mostly manual, wherein the langur is interacting directly (begging postures where langurs touch the humans) or indirectly (passive begging, no direct contact between langur and human) with human beings.

Our study has also looked into the factors that govern a ‘successful begging’ event. A successful begging event is one when the free-ranging Hanuman langur had received the food item that it was asking for, from the human being. Since the DG troop is female adult skewed, begging events are largely manifested by them, irrespective of its success. BGc was recorded as the most common form of begging gesture, however for making the event successful, BGpi was most efficient followed by BGe. Principal Component analysis further strengthen this observation. The mean and median number of successful begging events were 11 and 7, respectively, except one outlier with 65 successful begging events. Moreover, most of these begging behaviours were recorded in the evening session, in contrast to morning and evening slots. These observations can thus be attributed to the fact that the temple area receives larger human flux during religious holidays, and during specified visiting time-slots, thereby facilitating higher human-langur interactions in terms of food provisioning, resulting in a surge of session-specific begging behavior and the occurrence of the outlier, when most of the tourists would come to visit the temple.

Considering their immense deity value, tourists and pilgrims who frequently visit the temple of Dakshineswar, intentionally come in close contact to these langurs and offer food items to them. In our previous work, we found that the DG langur troop had a clear preference for bun (processed bread) which they can select from a given range of processed and unprocessed food items (Dasgupta et al., 2021). Therefore, these langurs acquire a generalised tendency to select the preferred food item and frequently reject the offered food until they received the desired one and continued to beg from the same person. However, most people did not give in to this persistent begging, resulting in a decrease in the correlation between successful begging and subsequent rounds of begging (Offer 1: r = 0.988; Offer 2: r = 0.325; Offer 3: *r* = 0.061). Furthermore, Random Forest Model and Artificial Neural Network also reiterated the pivotal role played by adult female langur, first begging sequence and provocation-initiated begging (BGpi) into contributing to the utmost success of the begging event.

Previous studies on ontogenetic and phylogenetic ritualization have suggested that gestures can be acquired through repeated use in an individual’s lifetime or inherited from previous generations (Call & Tomasso, 2007; Byrne et al., 2017). Here, our study on the Dakshineswar langur troop, which is heavily provisioned and comfortable in the presence of humans, found that gestural communication is learned over time through imitation of conspecifics as the correlation with successful begging events increased with age (Adult: r = 0.965; Subadult: r = 0.502; Juvenile: r = 0.067). Moreover, the reward received at the end of the begging action is acting as a reinforcement for this behaviour. These outcomes appear to be the result of ontogenetic ritualization as IGC seems to be acquired by this troop through repeated usage during it’s lifetime. Additionally, these successful begging events seem to be an effective foraging strategy for urban-adapted free-ranging species like the DG langurs who can acquire high-calorie processed food items as opposed to their natural plant-based, low-calorie diet. Interestingly, existing research literature points out that IGC is present among several NHPs including both catarrhine and platyrrhine lineage (Schell,2022). If IGC is present among most of the species in the Primate order, it might be possible that phylogenetic ritualization might be at play here. However, further studies across the species belonging to the Primate order will be required to make any concrete claims. However, considering the successful existence of such IGC which enhances the cooperative human-langur interactions, this could be a prospective area that should be further investigated by conservationists for the successful coexistence of free-ranging, homeless animals within human-modified ecosystem and promotes the establishment of a sustainable urban ecosystem.

## Conclusion

Our study provides the first systematic evidence of intentional gestural communication in free-ranging Hanuman langurs in Dakshineswar, West Bengal, India. This observation supports the idea that the evolution of language in humans may have been built on pre-existing communication abilities found in other non-human primates. To better understand the evolution of language in primates, more comprehensive studies across all lineages of primates, preferably in free-ranging contexts, are required. Additionally, research on the cognitive and neural mechanisms underlying language evolution and development is highly imperative. By studying the communication abilities of non-human primates, we can gain insights into the evolution of language and the development of language in humans. Simultaneously, such instances of intentional gestural communication in free-ranging urban animals promote cooperative human-animal interactions, further intensifying the notion of successful coexistence within urban ecosystems.

## Acknowledgments

DD and AB equally contributed to this work. DD, AD, SM, DB, RK, SK, ABh, and SG carried out the fieldwork. MP, DD, AD coded the entire data. AB, DD and MP carried out all the statistical analyses. AB ran the PCA, ANN, and RFM models to interpret the data. MP conceptualized the study, got grants to support the work, designed the fieldwork and supervised the work. DD, MP and AD drafted the manuscript. PB provided laboratory support to carry out the analysis. DD acknowledges the Prime Minister’s Research Fellowship, Ministry of Education, Govt of India for providing her fellowship. All the authors acknowledge Sourav Biswas for his digital art depicting all the begging postures.

